# Evaluating Stock Status and Biological Reference Points for a Data-Limited Brown Shrimp Fishery in the Gulf of California Data-Limited Assessment of the brown shrimp fishery in the Gulf of California

**DOI:** 10.1101/2025.06.07.658454

**Authors:** Francisco Arreguín-Sánchez, Carlos Iván Pérez-Quiñónez, Armando Hernandez-Lopez, Darío Chávez Herrera

## Abstract

The state of exploitation of the brown shrimp, *Penaeus californiensis*, fishery in the northeastern region of the Gulf of California was evaluated using landed catch and fishing efforts from 2010 to 2020. The annual catchability coefficient, 𝑞_𝑦_, was estimated through the Leslie model; the monthly biomass, 𝐵_𝑚_, was estimated from the catch per unit effort; 𝑈_𝑚_ via the relationship 𝐵_𝑚_ = 𝑈_𝑚_/𝑞_𝑦_; and through a biomass accumulation model throughout the fishing season, an approach to the biomass corresponding to the carrying capacity, 𝐵_∞,𝑦_, was obtained. The harvest rate, 𝐻𝑅, for each month and year was obtained and the annual maximum sustainable yield, 𝑀𝑆𝑌_𝑦_, derived as 𝐻𝑅_𝑀𝑆𝑌,𝑦_ = 0.50, which represents the target biological reference point, 𝐵𝑅𝑃_𝑡𝑔𝑡_. The decrease in population abundance allows for estimates of the recruitment rate, 𝜌_𝑦_, at the beginning of the fishing season and the survival rate, 𝑠_𝑦_, at the end of the fishing season, which represent the remaining reproductive population. From these estimates, a limit biological reference point was computed, described as 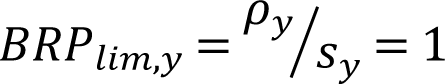, which corresponds to the population replacement level. With these estimates, the Kobe diagram was constructed, in which the biological reference points 𝐵𝑅𝑃_𝑡𝑔𝑡_ and 𝐵𝑅𝑃_𝑙𝑖𝑚,𝑦_ were incorporated. The diagnosis of the state of exploitation indicates, considering a precautionary criterion regarding recruitment, that the fishery has been operating at acceptable levels of sustainability, near the maximum production capacity.

## Introduction

Shrimp fisheries are the most important for Mexico, both economically and in terms of job creation, followed by tuna and sardine fisheries [1]. According to “daily landing notices” recorded by the National Commission of Fisheries and Aquaculture, CONAPESCA, for 2023, the total catch in the northeastern Gulf of California (Sonora and Sinaloa littorals) was 34,105 tons, with a landing value of 133 million dollars, corresponding to 67% of national production. The most important species in this region are blue shrimp (*Penaeus stylirostris*) (accounting for 74% of total catches), brown shrimp (*P. californiensis*) (21%), white shrimp (*P. vannamei*) (4%), and other shrimp species account for the remaining 1.1%.

The Instituto Mexicano de Investigación en Pesca y Acuacultura Sostenibles, IMIPAS, recognises five shrimp regions on the Mexican Pacific coast (Fig 1) and annually assesses the state of exploitation of the fish resources. The results of these assessments are reflected in the National Fisheries Charter, NFC [2], where management criteria for their application are also defined. For example, the most recent shrimp fishery report [2] indicates that the shrimp resources on the Pacific coast are exploited at their maximum sustainable yield, defined as a management strategy for the maintenance of a minimum reproductive biomass at the end of the fishing season and as management tactics, the control of fishing efforts, the establishment of a spatial temporal reproductive and growth ban, and the regulation of fishing gears. However, these guidelines do not specifically mention the state of exploitation by species, regions, or specific biological reference points, information that is relevant given the diversity of species, habitats, and different responses to changes in climatic patterns, or specific interest of the different fishing communities in accessing fishery certification processes.

**Fig 1.**
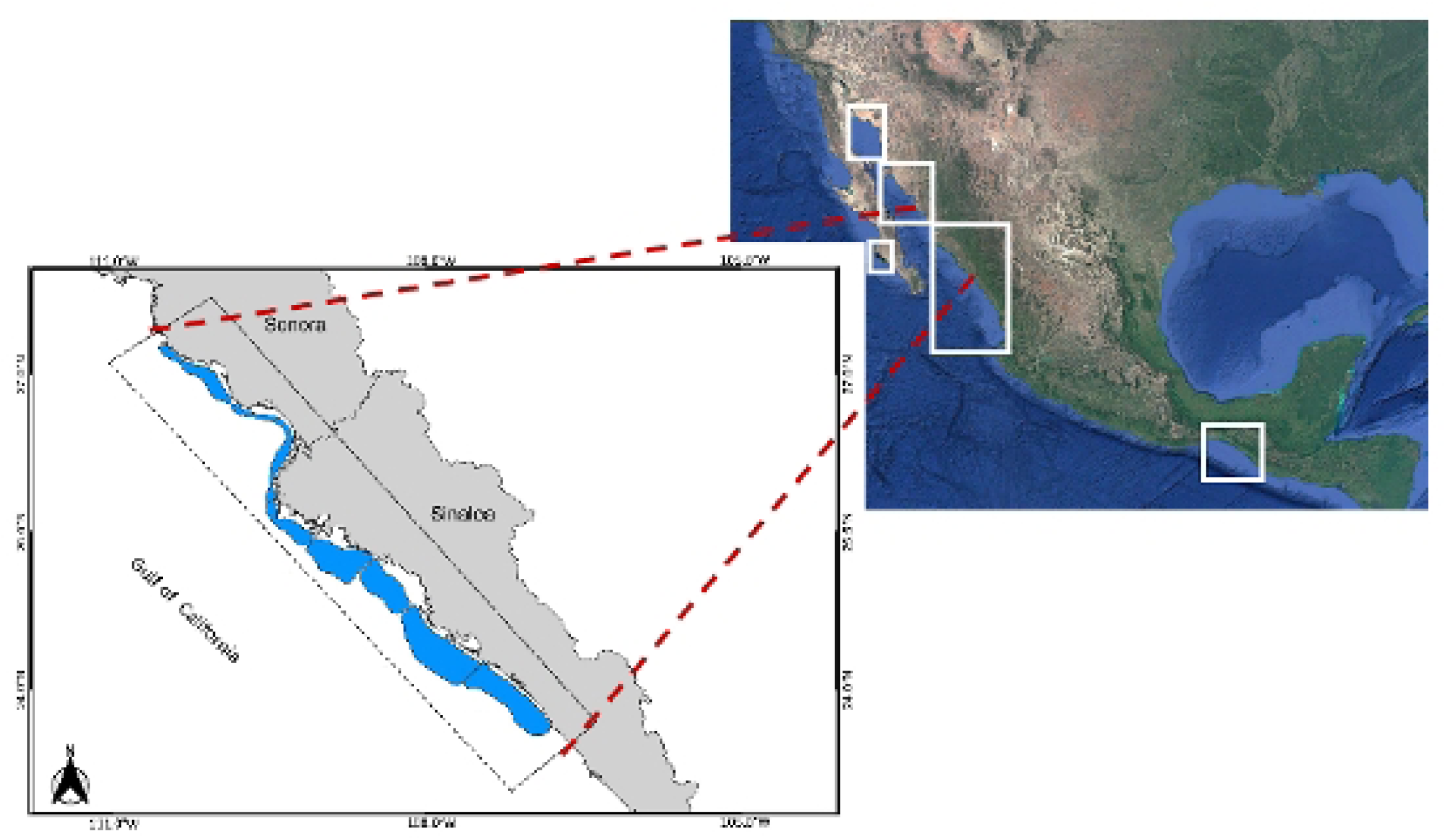
Main shrimp fishing areas on the Mexican Pacific coast and study region [33]. White squares show Pacific fishing areas. The inset on the left shows the study region for the analysis of the brown shrimp fishery.

A review of the scientific literature reflects only 19 contributions to the assessment of the state of exploitation of shrimp fisheries on the Mexican Pacific coast, covering all species from 1949 to 2021 (Table 1). Specifically, the brown shrimp fishery is mentioned in only 13 of them; the most recent works correspond to the periods 2000 to 2014 for the southern Gulf of California [3]; from 1976 to 1998 for the Sonora and Sinaloa region [4]; and for the 1995/1996 fishing season for the northern Gulf of California [5].

**Table 1.**
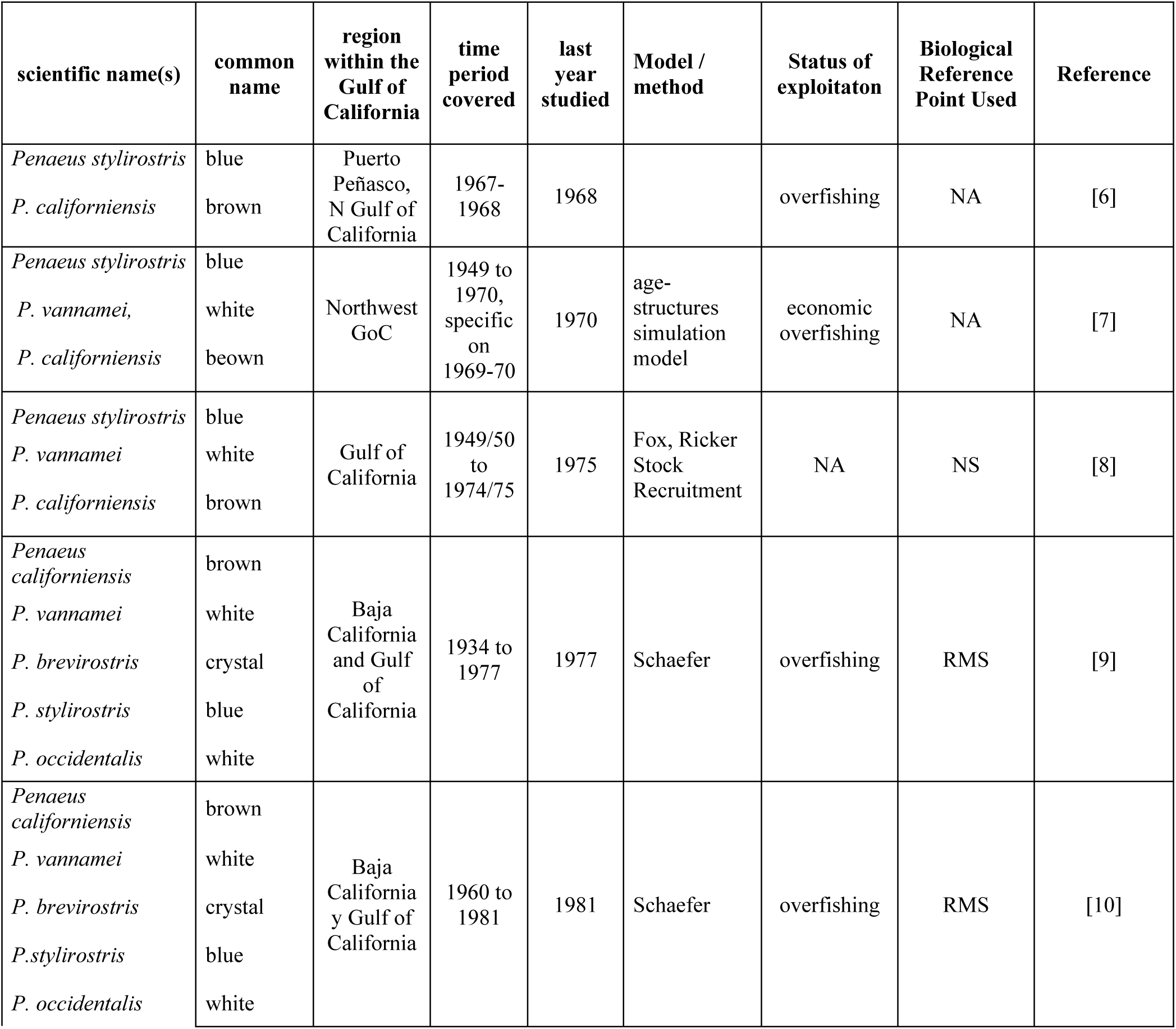
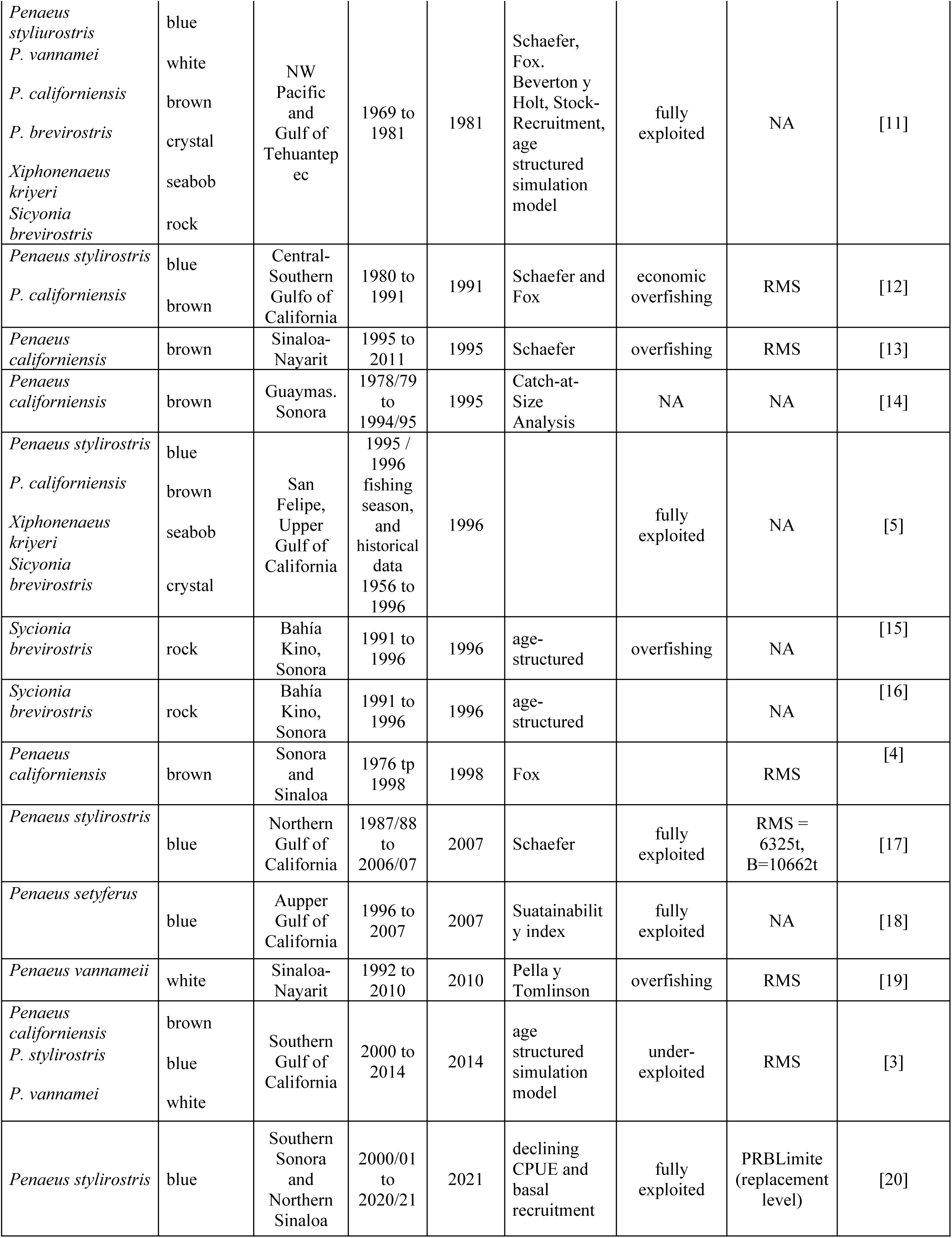
Reports in the literature focused on stock assessment and the state of exploitation of shrimp resources in the Gulf of California.

On the other hand, IMIPAS conducts annual studies on the reproductive status of the most important species to determine the start and end dates of a reproductive ban [i.e. 21], as well as assessments of the state of exploitation of shrimp resources, and the results are included in the NFC [2]; however, in this case, these documents remain internal technical reports. In general, most scientific contributions use the maximum sustainable yield as a reference point, and only one study defines a limit reference point for blue shrimp [20].

There are, of course, a significant number of scientific contributions to the knowledge of shrimp resources in the Gulf of California; for example, for brown shrimp, contributions to population dynamics include aspects of biometric relationships [22], individual growth [23, 24], natural mortality [25], catchability [26], and spatial distribution [27], as well as those related to environmental effects [28, 29, 30, 31, 32].

A relevant aspect from a methodological point of view is that several assessments of the brown shrimp fishery, including those that give rise to the NFC, are based on the application of the dynamic biomass model, probably due to the limited information available beyond catch and effort records. In this sense, the conceptual biological bases of these models are not applicable to species with annual life cycles. According to Hilborn and Walters [34], the Schaefer model [35, 36] is described, in its discrete form, by the following equation:

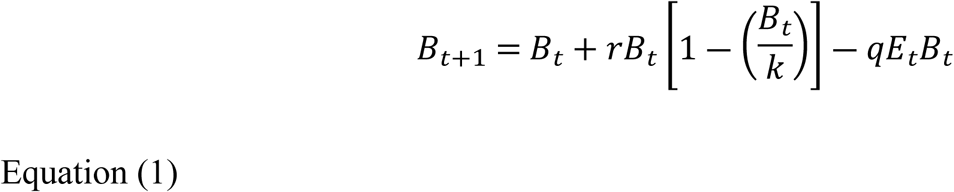

where 𝐵 represents biomass; 𝑡 represents the unit of time, typically annual to include complete reproduction periods, and therefore recruitment; 𝑟 represents the intrinsic rate of population growth;𝑘 the carrying capacity defined by the environment; and 𝑞 represents the catchability coefficient; with the three parameters assumed to be constant for the analysed period. Additionally, 𝐸 represents the fishing effort, and 𝑞𝐸_𝑡_𝐵_𝑡_ represents the catch at time 𝑡. The reason why the model expressed by Equation (1) is not applicable to annual species is that when 𝐵_𝑡+1_ ≠ 0, then 𝐵_𝑡_ = 0, and vice versa; consequently, Equation (1) has no solution. On the other hand, if values of year 𝑡 < 1 are considered, then 𝑟 and 𝑘 cannot be assumed to be constant since they would change intra-annually, year Δ𝑡 < 1 (for example, seasonal changes).

An additional problem in analysing fisheries of annual species, such as penaeid shrimp, is the limited availability of data that allows for a clear notion of the fishery’s evolution and the state of exploitation of the brown shrimp stock over time, allowing for the definition of biological reference points, 𝐵𝑅𝑃𝑠, to guide management. In this context, the objectives of this work were i) to assess the state of exploitation of the brown shrimp resource in the northeastern Gulf of California (Fig 1) using only catch and fishing effort data; ii) to define the fishery’s evolution over the last two decades on the basis of the target biological reference points, 𝐵𝑅𝑃_𝑡𝑔𝑡_, and limit 𝐵𝑅𝑃_𝐿𝑖𝑚_; and iii) to suggest some aspects directly applicable to resource management.

## Matherials and methods

### Data Sources and Organisation

Information on catch (live weight, tons) and fishing effort (effective fishing days) for small-scale and industrial fleets, recorded in daily landing notices, was available through CONAPESCA for the period 2010 to 2021. The small-scale fleet, 𝑠𝑠𝑓, uses various types of fishing gear (“suripera”, trawl and cast nets), and the industrial fleet, 𝑖𝑛𝑑𝑓, operates with bottom trawl nets. In the absence of more detailed temporal quantitative information, the participation of both fleets is assumed: each fleet has its own fishing power, and both operate simultaneously on the same resource in time and space. Thus, if the fishing power between fleets, 𝑖 and 𝑗, is similar, the relationship between the catch per unit effort, 𝑈, per month, 𝑚, should be 𝑈_𝑚,𝑖_ = 𝑏𝑈_𝑚,𝑗_, where 𝑏 = 1; however, if 𝑏 ≠ 1, the value of 𝑏 corresponds to the conversion factor of the fishing power of one fleet relative to the other.

### State of Exploitation

In general terms, 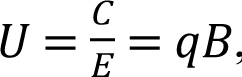, where 𝐵 represents biomass and 𝑞 represents the catchability coefficient. To estimate 𝑞, the Leslie model [37, 34] is used each year, assuming a closed population, a criterion that can be accepted for a species with annual longevity. This model is represented by the following relationship:

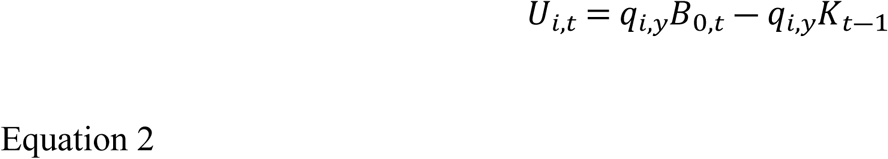

where 𝑞_𝑖_ is the catchability coefficient for fleet 𝑖, assumed constant for year 𝑦; 𝑈_𝑖,𝑡_ is the catch per unit effort of fleet 𝑖 at time 𝑡, in our case 𝑡 = one month; 𝑈_0,𝑡_ is the intercept (𝑞_𝑖,𝑦_𝐵_0,𝑡_) in Equation 2; and 𝐵_0,𝑖,𝑡_ is the population size available to fleet 𝑖 just before the first observed data point and can be estimated as 𝐵_0,𝑡_ = 𝑈_0_/𝑞_𝑖,𝑦_. 𝐾_𝑡―1_ is the cumulative catch at time 𝑡 ―1. Similarly, the population biomass in month 𝑚 is as follows:

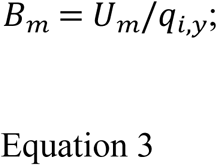

and the population size during the fishing season is:

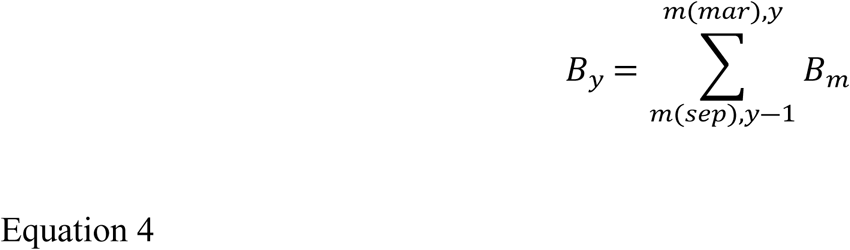

Thus, the state of exploitation of the population can be determined through the harvest rate, expressed as 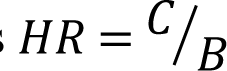.

In the case of the brown shrimp fishery, the fishing season starts in September and ends in March. The closed season is defined to protect the reproductive biomass present at the end of the season, as well as to ensure that growth overfishing does not occur by August, when recruits join the fishery. Therefore, as an annual species, 𝐵_0,𝑡_ corresponds to the biomass in August, and 𝐵_𝑠𝑒𝑝_ corresponds to the biomass of recruits available for fishing in September; and consequently, 𝐵_𝑚𝑎𝑟_ corresponds to the reproductive biomass, in March, at the end of the fishing season.

On the other hand, although the theoretical assumption of the dynamic biomass model does not correspond to species with annual longevity, it is possible to represent a biomass accumulation model throughout the year, such that the parameters 𝑟 and 𝑘 of the Schaefer model [35, 36] (intrinsic growth rate and carrying capacity, respectively) can be replaced by the biomass accumulation rate 𝛿_𝑦_ throughout the year and the population size in the year, as an approximation to 𝐵_∞,𝑦_. This model is represented as:

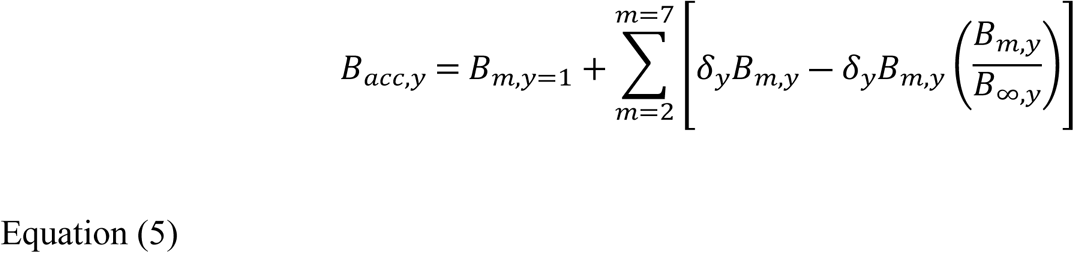

where for the last time-period 𝑚, 𝐵_𝑎𝑐𝑐,𝑦_ = 𝐵_∞_.

### Biological Reference Points

According to Gulland [38], for a stable population in a stable environment, the maximum sustainable yield, 𝑀𝑆𝑌, is defined by the harvest rate 𝐻𝑅_𝑀𝑆𝑌_ = 0.5. When the population state relative to the value 𝑘 (Equation 1) is at a level 𝑘/2, corresponding to the population state where the maximum biomass production capacity is reached, it is defined as the target biological reference point, 𝐵𝑅𝑃_𝑡𝑔𝑡_; that is, when 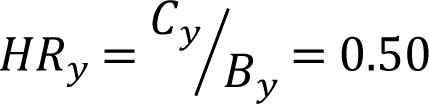. Similarly, an analogous calculation could be made for each month to determine the monthly harvest rate using the available biomass in the month, 𝐵_𝑚_.

On the other hand, Arreguín-Sánchez et al. [20]) proposed a limit biological reference point, 𝐵𝑅𝑃_𝑙𝑖𝑚_, as the population replacement level; that is, when the adult population level is equal to the recruit population level, it is necessary to maintain a population with constant density [39, 40, 41, 42]. To define 𝐵𝑅𝑃_𝑙𝑖𝑚_, the annual recruitment rate 𝜌_𝑦_ at the start of the fishing season (September of year 𝑦) is considered relative to the adult population that gave rise to it the previous year (March of year 𝑦 ―1), indicated by the survival rate, 𝑠_𝑦_, represented as:
the recruitment rate

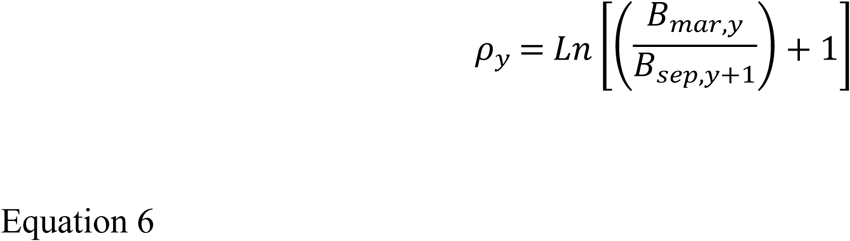

and survival rate

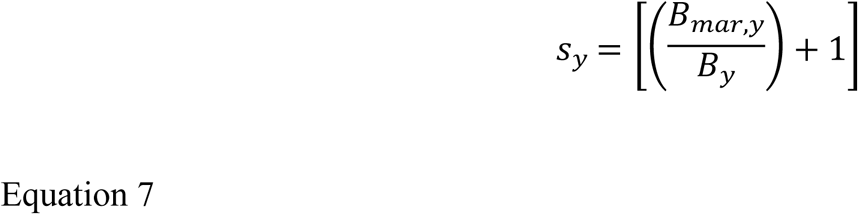

Thus, the replacement level corresponding to 𝐵𝑅𝑃_𝑙𝑖𝑚,𝑦_ is defined when 𝜌_𝑦_ = 𝑠_𝑦_; that is,

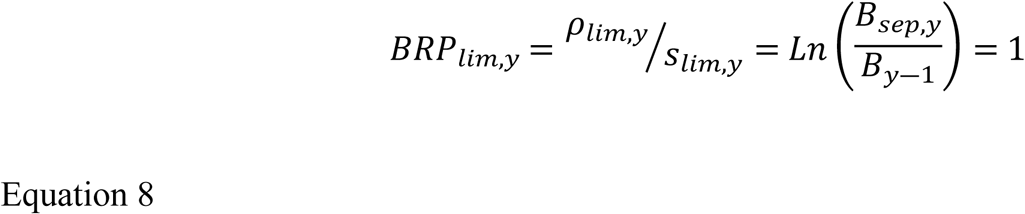

The above equality refers to the condition under which the survival and recruitment rates are similar, representing the population replacement level [43], and equivalent to the basal recruitment level [44]. To obtain 𝐵𝑅𝑃_𝐿𝑖𝑚,𝑦_, a graph of 𝜌_𝑦_ relative to 𝑠_𝑦_ was constructed, where the intersection with the bisector where 𝜌_𝑙𝑖𝑚,𝑦_ = 𝑠_𝑙𝑖𝑚,𝑦_ identifies the limit levels for 𝜌_𝑙𝑖𝑚,𝑦_ and 𝑠_𝑙𝑖𝑚,𝑦_.

As noted by Shepherd [39], there is no single replacement level given the interannual variability of populations, which is especially noticeable in species with annual longevity; therefore, a different replacement level is expected for each annual combination of recruits and adults.

### Fishery Evolution and Management Orientation

For the Kobe diagram, the ratios 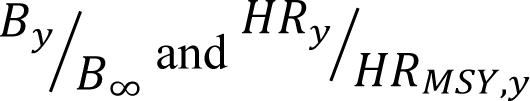 were used, assuming that the maximum sustainable yield, 𝑀𝑆𝑌, refers to a harvest rate 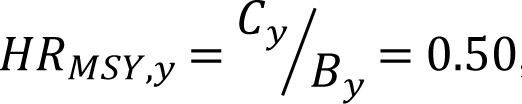, representing the target biological reference points, 𝐵𝑅𝑃_𝑡𝑔𝑡_. With these estimates for each year, the evolution of the state of exploitation can be observed relative to 𝐵𝑅𝑃_𝑡𝑔𝑡_, and 𝑃𝑅𝐵_𝑙𝑖𝑚_. Thus, depending on the fishery’s condition, both historically and currently, the possible application of these 𝐵𝑅𝑃𝑠 in daily management practices can be discussed, with the NFC [2] as a reference framework.

## Results

### Data, Fishing Power, and Catchability

The fleet operation records were organised by fishing season and month, and their trends, U_m,y_, clearly decreased as the fishing season progressed (Fig 2). With respect to the fishing effort application pattern, a decreasing pattern was also observed for the 𝑖𝑛𝑑𝑓 fleet, following abundance changes, whereas the 𝑠𝑠𝑓 fleet operation pattern shows greater variability.

**Fig 2.**
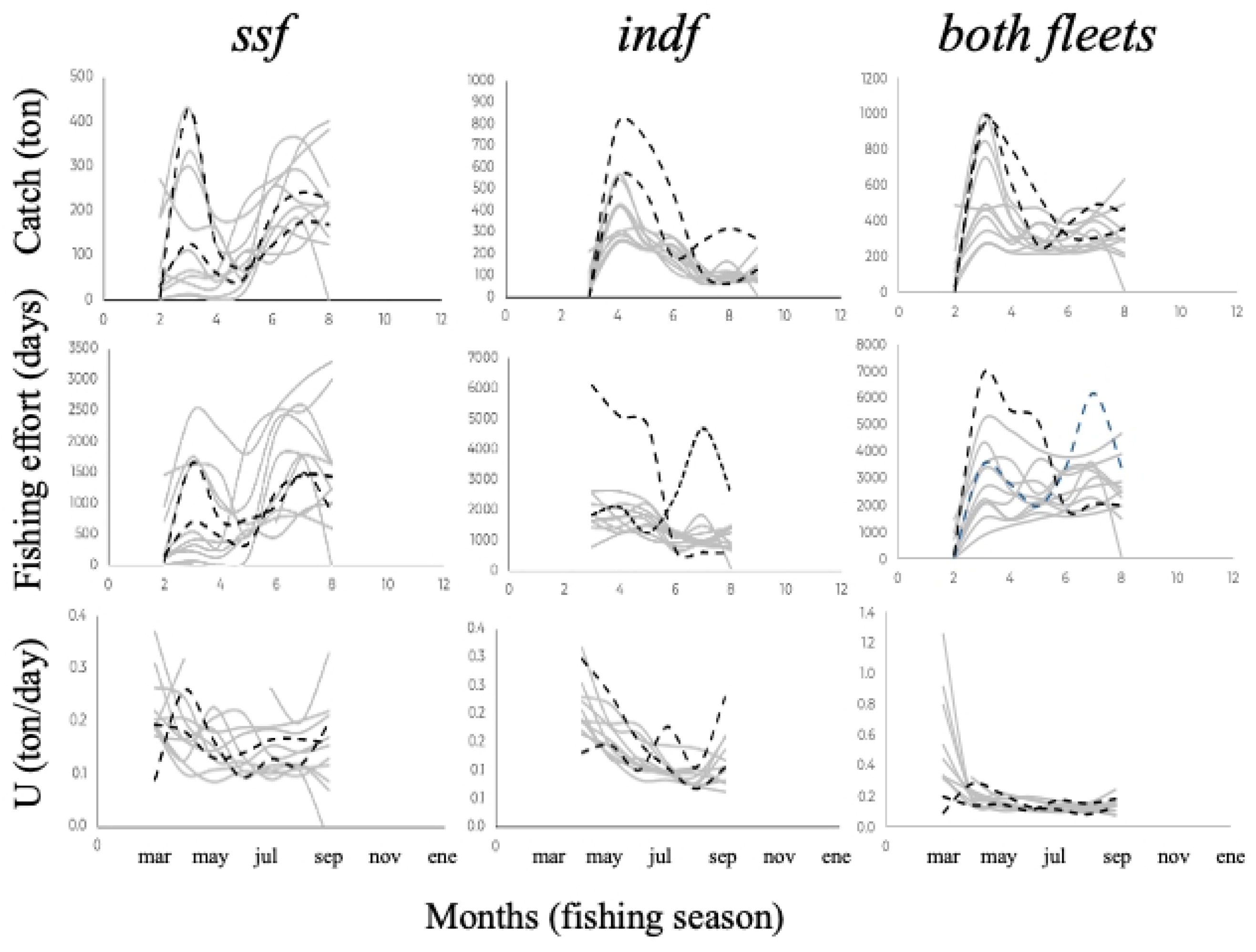
Trends in the catch, fishing effort, and catch per unit effort by fleet and fishing season for the brown shrimp fishery in the northwest Gulf of California from 2010 to 2021. The grey lines represent different fishing seasons, whereas the black dashed lines correspond to the COVID-19 pandemic years. 𝑠𝑠𝑓 = small scale fleet; 𝑖𝑛𝑑𝑓 = industrial fleet.

Regarding the fishing power of the fleets, Fig 3 shows two different patterns: the first pattern was present in 8 fishing seasons, where the fishing power of the 𝑠𝑠𝑓 fleet exceeded that of the 𝑖𝑛𝑑𝑓 fleet by a factor of 3, whereas the second pattern was reversed during 3 fishing seasons (2011/12, 2012/13, and 2014/15), in which the fishing power of the 𝑖𝑛𝑑𝑓 fleet surpassed that of the 𝑠𝑠𝑓 fleet by a factor of 2.3.

**Fig 3.**
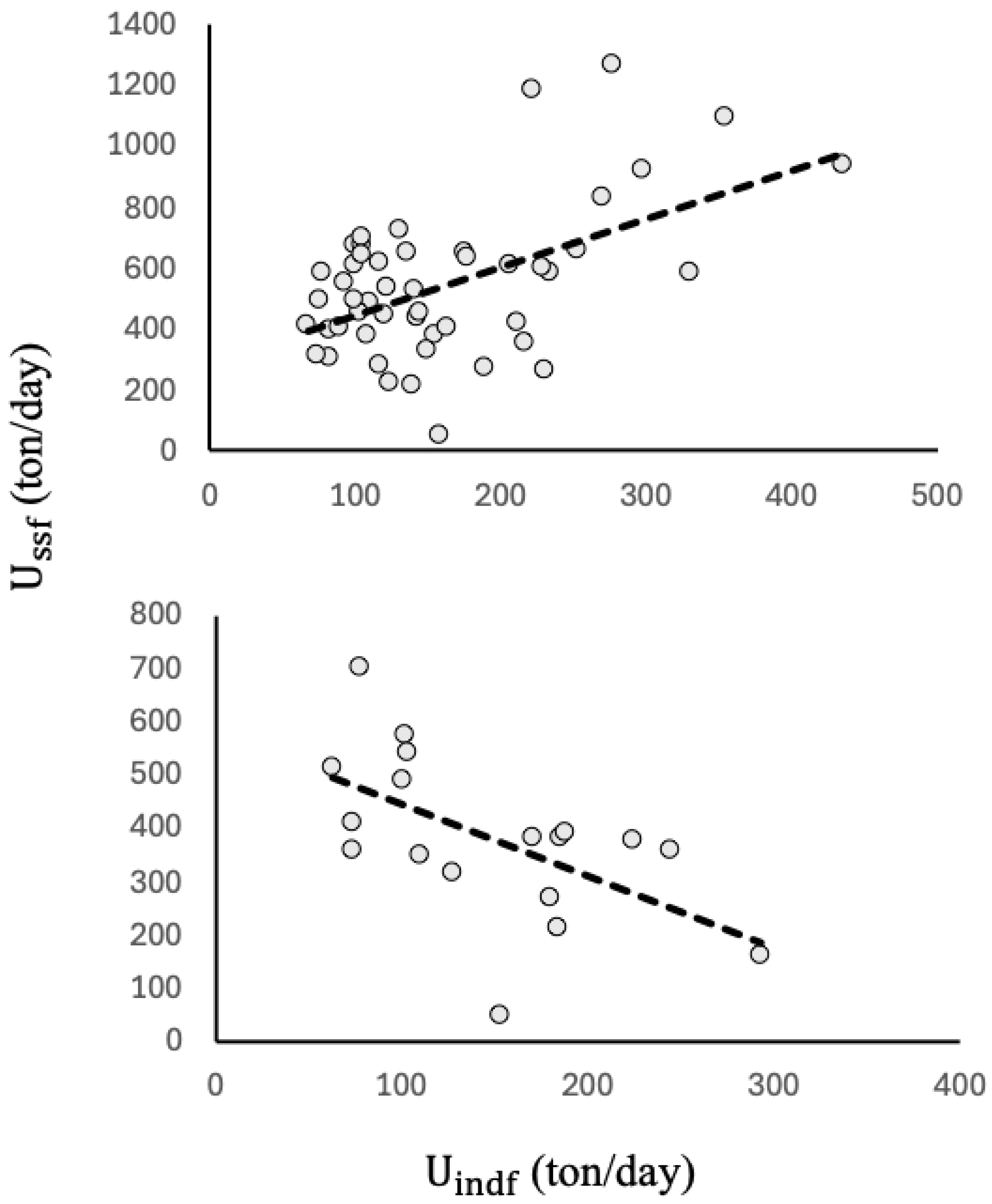
Comparison of fishing power between fleets of the brown shrimp fishery in the northwest Gulf of California. The ssf fleet showed a fishing power superior to that of the indf fleet by a factor of 3 in 8 fishing seasons; the indf fleet had greater fishing power than the ssf fleet by a factor of 2.3 in 3 fishing seasons (S 1 Tables).

To identify the state of exploitation of the resource, the catchability coefficient, 𝑞 (Equation 2), was initially estimated, representing the probability that a unit of population is retained by a unit of effort [45]. In our case study, although the unit of effort for both fleets is the effective day, it is evident that the fishing day of the 𝑠𝑠𝑓 fleet is different from the effective fishing day of the 𝑖𝑛𝑑𝑓 fleet. For this reason, the catchability coefficient for each year and fleet 𝑞_𝑖,𝑦_ was estimated, as shown in Table 2. Generally, the catchability for the 𝑠𝑠𝑓 fleet was slightly greater than that for the 𝑖𝑛𝑑𝑓 fleet, with the following average values: 𝑞_𝑠𝑠𝑓,𝑦_ = 0.00021 (𝑐𝑣 = 40%) and 𝑞_𝑖𝑛𝑑𝑓,𝑦_ = 0.00015, (𝑐𝑣 = 43%).

**Table 2.**
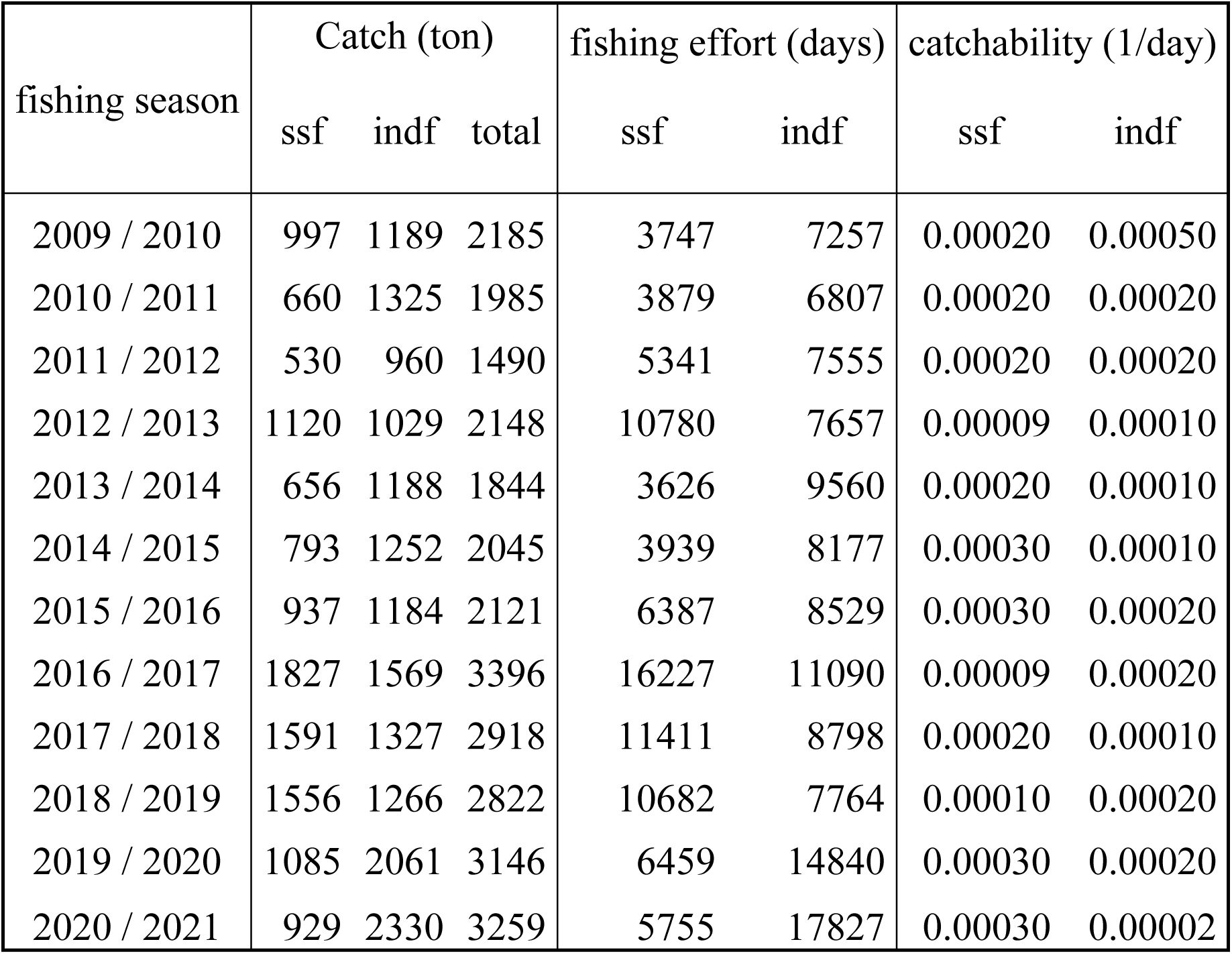
Annual records of catch and fishing efforts, and catchability estimates by fleet, all by season, for the brown shrimp fishery in the northwest Gulf of California.

| fishing season | Catch (ton) |  |  | fishing effort (days) |  | catchability (1/day) |  |
| --- | --- | --- | --- | --- | --- | --- | --- |
|  | ssf | indf | total | ssf | indf | ssf | indf |
| 2009 / 2010 | 997 | 1189 | 2185 | 3747 | 7257 | 0.00020 | 0.00050 |
| 2010 / 2011 | 660 | 1325 | 1985 | 3879 | 6807 | 0.00020 | 0.00020 |
| 2011 / 2012 | 530 | 960 | 1490 | 5341 | 7555 | 0.00020 | 0.00020 |
| 2012 / 2013 | 1120 | 1029 | 2148 | 10780 | 7657 | 0.00009 | 0.00010 |
| 2013 / 2014 | 656 | 1188 | 1844 | 3626 | 9560 | 0.00020 | 0.00010 |
| 2014 / 2015 | 793 | 1252 | 2045 | 3939 | 8177 | 0.00030 | 0.00010 |
| 2015 / 2016 | 937 | 1184 | 2121 | 6387 | 8529 | 0.00030 | 0.00020 |
| 2016 / 2017 | 1827 | 1569 | 3396 | 16227 | 11090 | 0.00009 | 0.00020 |
| 2017 / 2018 | 1591 | 1327 | 2918 | 11411 | 8798 | 0.00020 | 0.00010 |
| 2018 / 2019 | 1556 | 1266 | 2822 | 10682 | 7764 | 0.00010 | 0.00020 |
| 2019 / 2020 | 1085 | 2061 | 3146 | 6459 | 14840 | 0.00030 | 0.00020 |
| 2020 / 2021 | 929 | 2330 | 3259 | 5755 | 17827 | 0.00030 | 0.00002 |

### Biomass Estimates and State of Exploitation

Once 𝑞 was estimated, and on the basis of Equation 3, the available biomass estimates for fleet 𝑖, month 𝑚, and fishing season 𝑦, 𝐵_𝑖,𝑚,𝑦_, was obtained, as well as the monthly population biomass as 𝐵_𝑚,𝑦_ = 𝐵_𝑖,𝑚,𝑦_ + 𝐵_𝑗,𝑚,𝑦_ and the total annual biomass according to Equation 4. The total biomass patterns by season are shown in Fig 4, and the estimated values by fleet, month, and year are presented in S 2 Tables.

**Fig 4.**
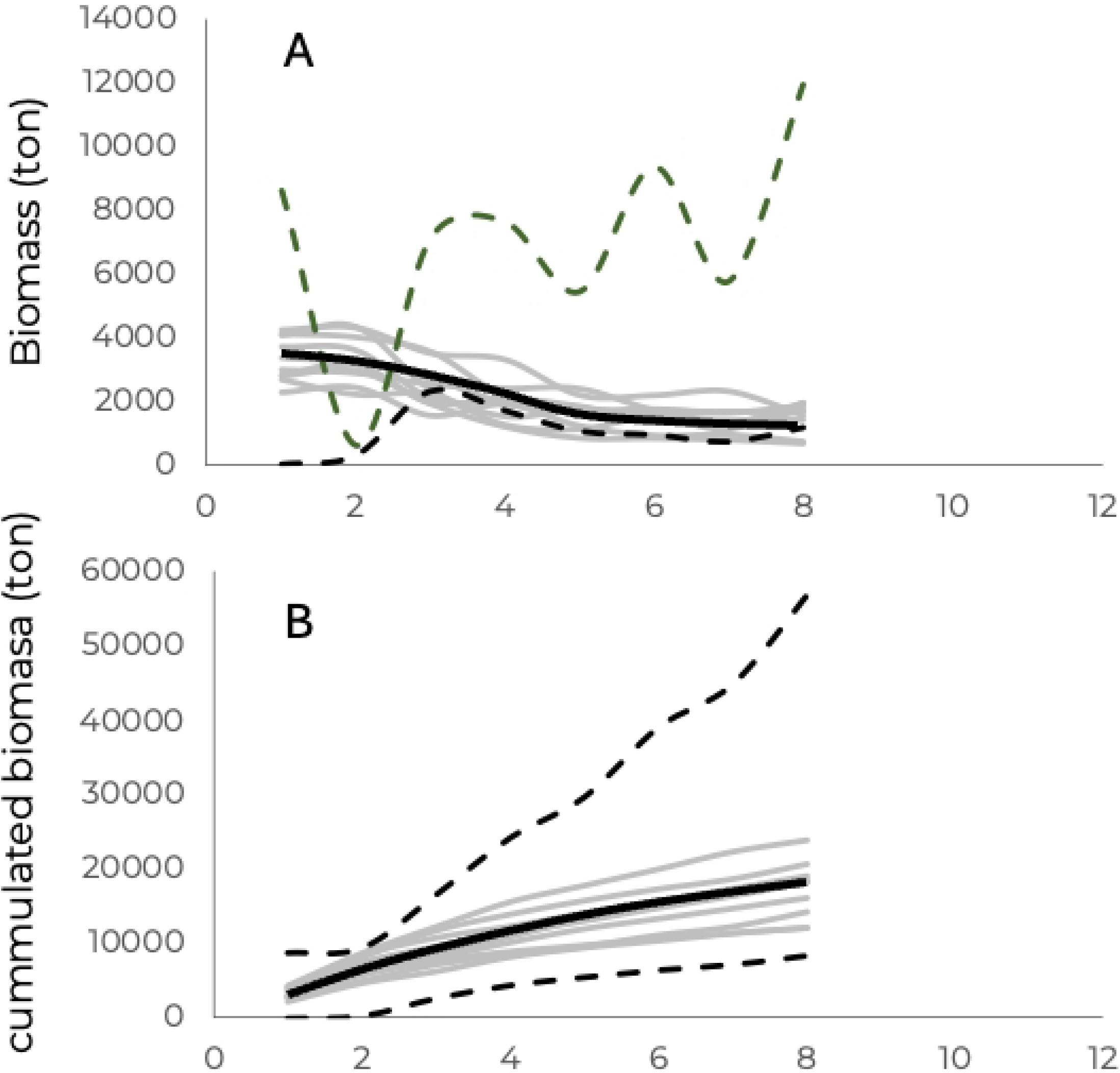
Trends in the biomass of the brown shrimp. *P. californiensis* **population in the northwest Gulf of California.** A) biomass decay and B) accumulated biomass throughout each fishing season. The dashed lines represent the years with the presence of the COVID-19 pandemic; the thick continuous lines represent the average patterns.

In addition, the estimation of 𝐵_∞,𝑦_ was carried out by applying the biomass accumulation model (Equation 5), where the biomass accumulation rate for each year, 𝛿_𝑦_, was also obtained. The resulting values in each case are shown in Table 3.

**Table 3.** Annual estimation of the parameters 𝛅_𝐲_ and 𝐁_∞,𝐲_ of the biomass accumulation model (**Equation 5**); and harvest rates: total, 𝐇𝐑_𝐲_; average, 𝐇𝐑_𝐚𝐯,𝐲_; weighted average, 𝐇𝐑_𝐰,𝐦𝐞𝐚𝐧,𝐲_; and survival, 𝐬_𝐲_; and recruitment 𝛒_𝐲_, rates.

| fishing season | $\delta_y$ | $B_{\infty,y}$ | $B_{cum}$ | Harvest rate | | | $s_y$ | $\rho_y$ |
| --- | --- | --- | --- | --- | --- | --- | --- | --- |
| | | | | $HR_{tot}$ | $HR_{av,y}$ | $HR_{w,mean,y}$ | | |
| 2009 / 2010 | 0.563 | 15980 | 14,187 | 0.154 | 0.369 | 0.265 | 0.1300 |  |
| 2010 / 2011 | 0.517 | 18239 | 16,227 | 0.122 | 0.175 | 0.175 | 0.0846 | 0.5200 |
| 2011 / 2012 | 0.669 | 13803 | 13,134 | 0.124 | 0.215 | 0.208 | 0.0500 | 0.4572 |
| 2012 / 2013 | 0.461 | 27444 | 23,772 | 0.090 | 0.144 | 0.138 | 0.1096 | 0.1300 |
| 2013 / 2014 | 0.504 | 15257 | 13,688 | 0.101 | 0.138 | 0.145 | 0.0911 | 0.2475 |
| 2014 / 2015 | 0.497 | 19607 | 18,249 | 0.112 | 0.161 | 0.156 | 0.0906 | 0.3596 |
| 2015 / 2016 | 0.687 | 12475 | 12,175 | 0.174 | 0.305 | 0.349 | 0.0500 | 0.4776 |
| 2016 / 2017 | 0.753 | 26628 | 19,080 | 0.178 | 0.268 | 0.276 | 0.0847 | 0.1600 |
| 2017 / 2018 | 0.446 | 21712 | 18,436 | 0.158 | 0.231 | 0.211 | 0.0938 | 0.3720 |
| 2018 / 2019 | 0.553 | 22286 | 20,599 | 0.137 | 0.190 | 0.208 | 0.0916 | 0.3640 |
| 2019 / 2020 | 0.72 | 9125 | 8,301 | 0.379 | 0.368 | 0.420 | 0.1332 | 0.6099 |
| 2020 / 2021 | 0.294 | 150152 | 56,782 | 0.057 | 0.160 | 0.090 |  | 0.1282 |

Additionally, the harvest rate, 𝐻𝑅, was estimated through the relationship 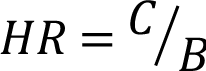; thus, the values for each month and fishing season were obtained. Three approximations of the annual harvest rate were obtained: the total harvest rate, 𝐻𝑅_*tot*_, represented for each year as 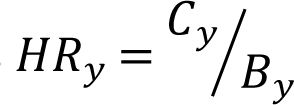; the average harvest rate for each year, which is based on the monthly harvest rates, 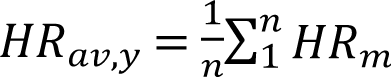; and the weighted average harvest rate, which uses 𝐻𝑅_𝑚_ and a weighting factor 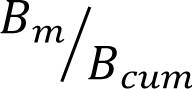, such that 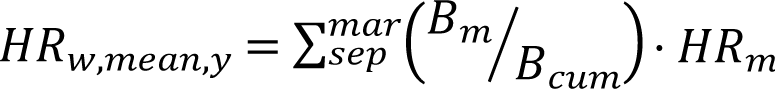. The estimated harvest rates for each year are shown in Table 3.

### The Limit Biological Reference Points

The recruitment rates, 𝜌_𝑦_, and survival rates, 𝑠_𝑦_, were estimated according to Equations 6 and 7, respectively (Table 3), and the limit biological reference point was estimated through Fig 5, corresponding to a value of 𝐵𝑅𝑃_𝑙𝑖𝑚,𝑦_ = 0.13 (Fig 5).

**Fig 5.**
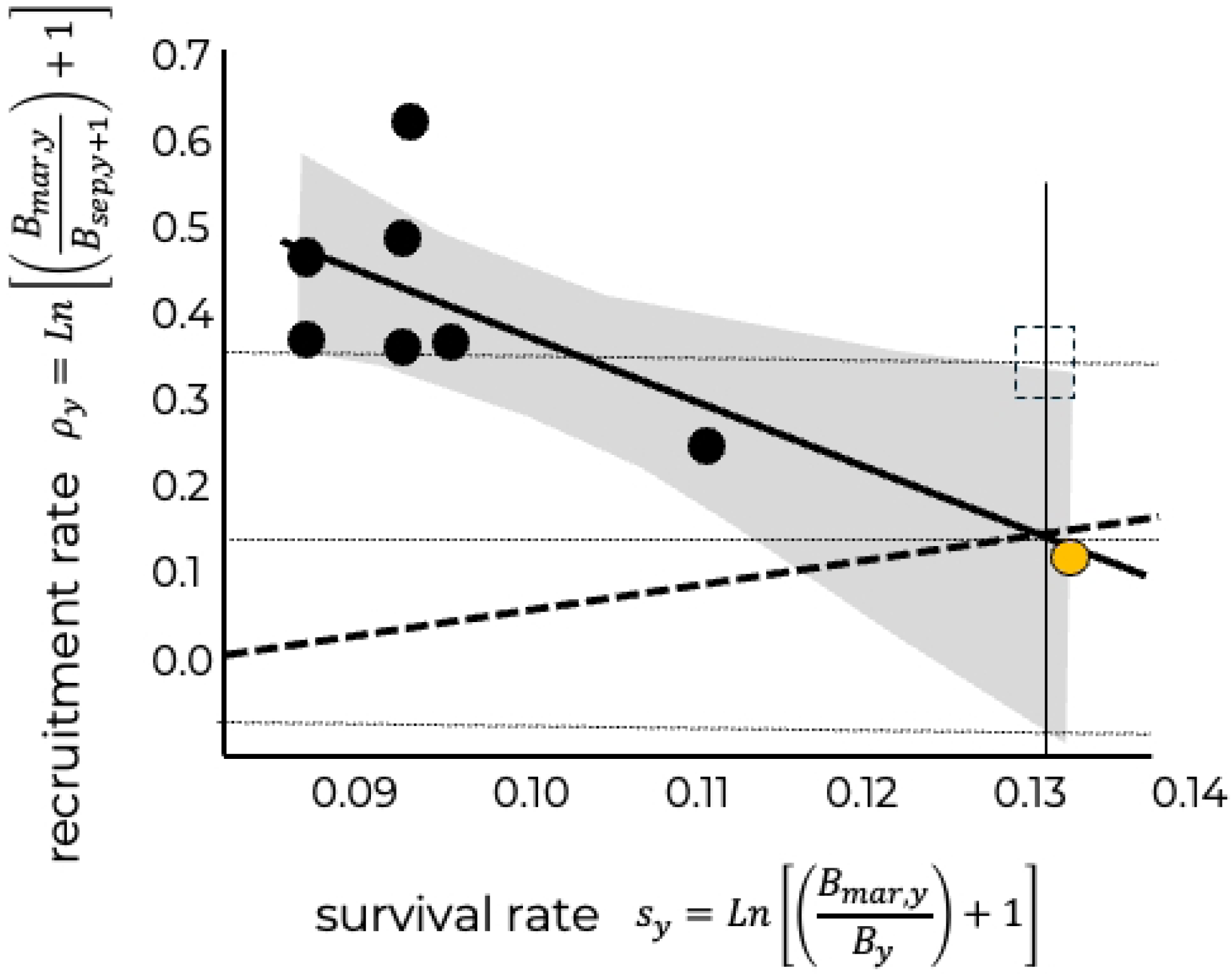
Identification of 𝐵𝑅𝑃_𝑙𝑖𝑚,𝑦_ for the *P. californiensis* brown shrimp fishery in the northwest Gulf of California. The 𝐵𝑅𝑃_𝑙𝑖𝑚,𝑦_ is given by the intersection point between the bisector (thick black dashed line) and the trend line of the relationship between the survival rate, 𝑠_𝑦_, and recruitment, 𝜌_𝑦_(continuous black line), where 𝑠_𝑦_ = 𝜌_𝑦_ = 0.13. The grey area corresponds to the 95% confidence interval of the regression, and the box with a dashed line indicates the limit recruitment rate level in a precautionary context, where 𝜌_𝑦_ = 0.38. The yellow point corresponds to the fishing seasons with the presence of the COVID-19 pandemic (2019/20 and 2020/21).

### Fishery Evolution and Kobe Diagram

The history of the interannual variation in the observed yields, 𝐶_𝑦_, of brown shrimp relative to the catch corresponding to the annual maximum sustainable yield, represented as 0.5 · 𝐵_∞,𝑦_, is shown in Fig 6. Note that for all the years of the studied period, 𝐶_𝑦_ < 0.5 · 𝐵_∞,𝑦_.

**Fig 6.**
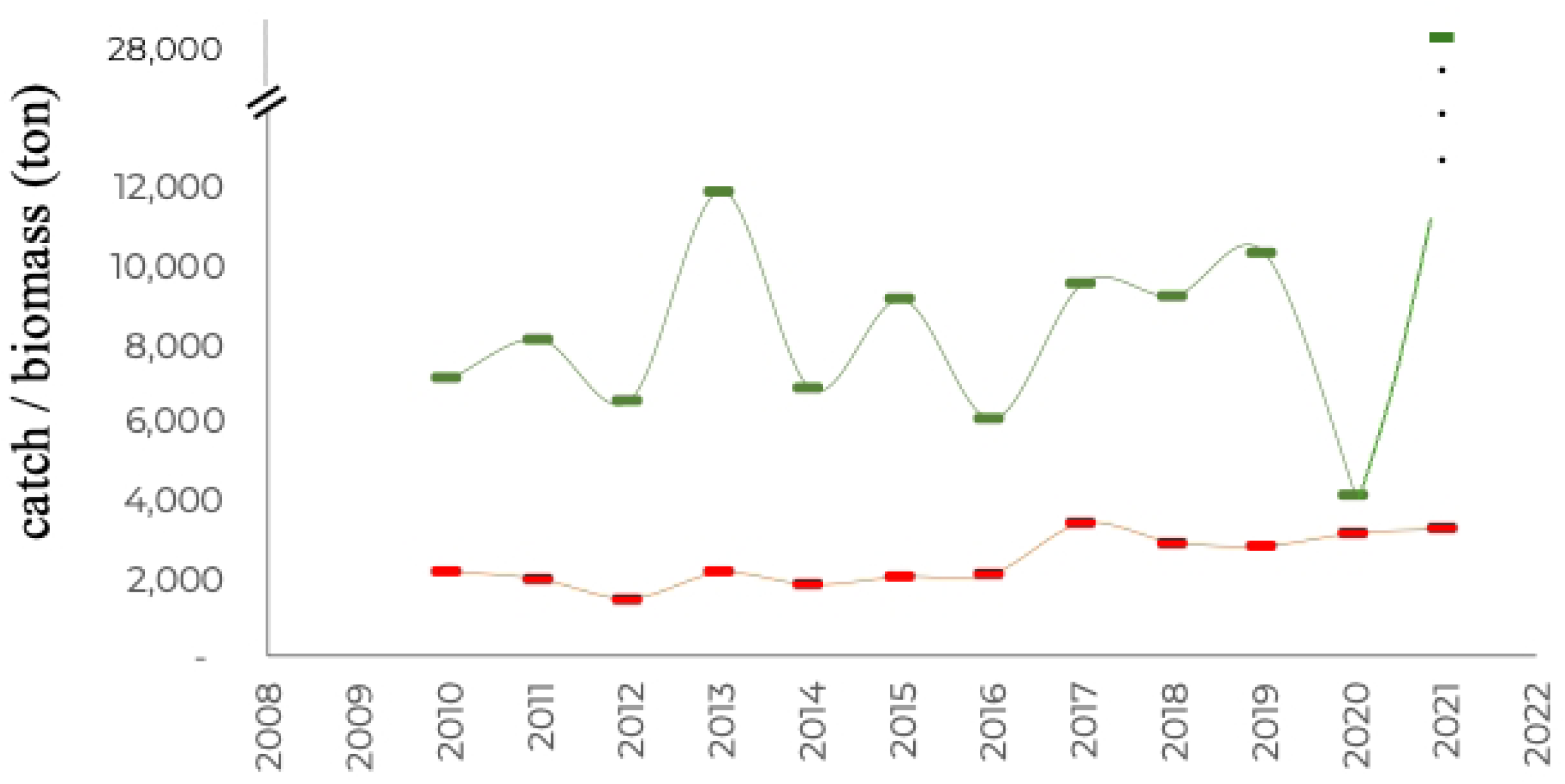
Historical evolution of the interannual variation of observed catches (red line) and the population level corresponding to 𝟎.𝟓 · 𝑩_∞,𝒚_ (green line). On the other hand, according to the conventional interpretation, the Kobe diagram (Fig 7) indicates that the brown shrimp fishery is being exploited at levels below the harvest rate corresponding to 𝑀𝑆𝑌, indicating that there is no overfishing; however, the ratio 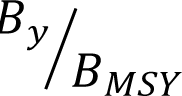 suggests some degree of depletion. Nevertheless, the fishery has remained stable throughout almost the entire analysed period, with values within the range 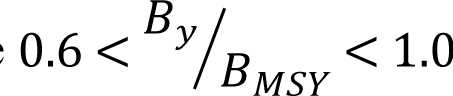.

**Fig 7.**
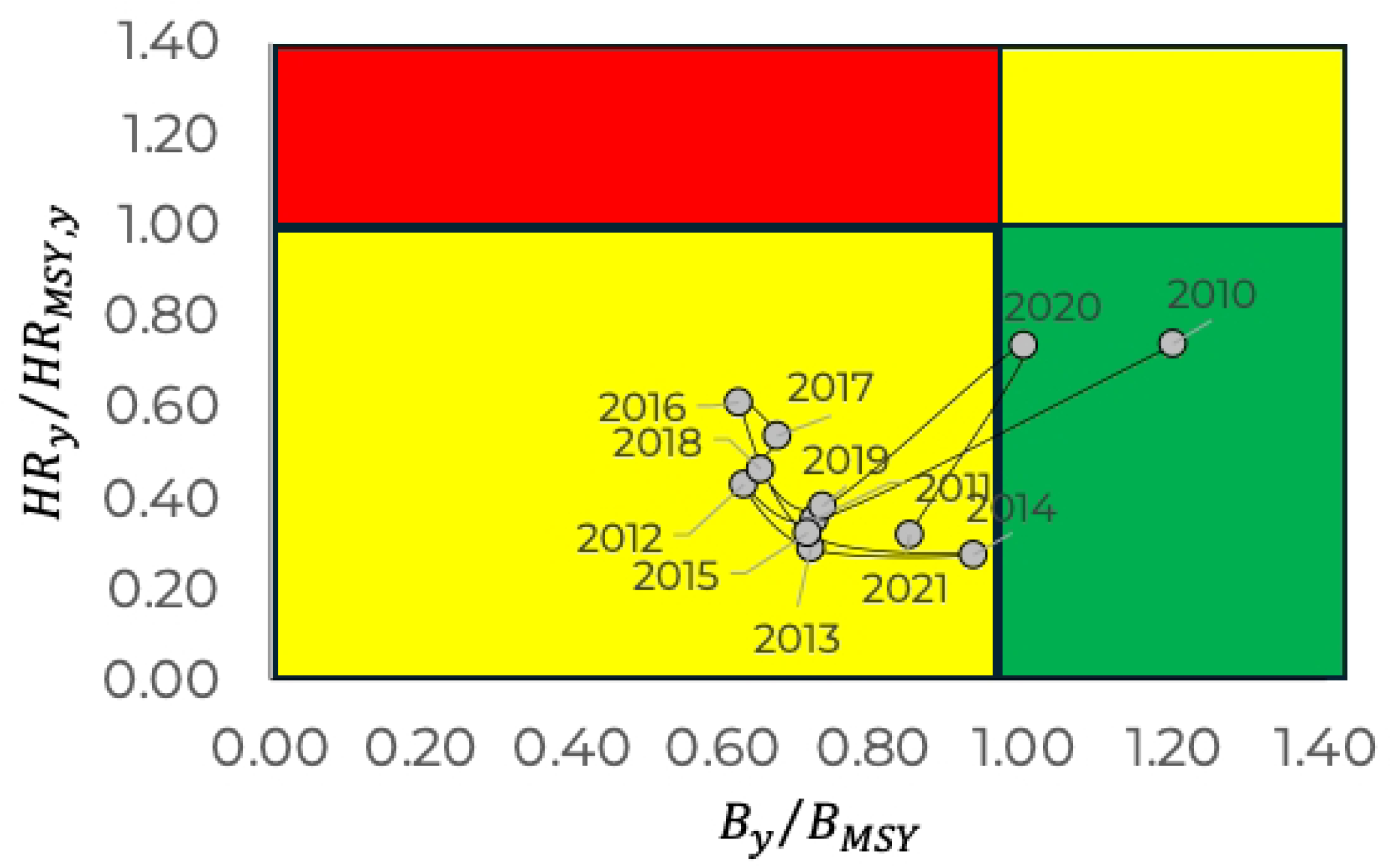
Kobe diagram for the *P. californiensis* brown shrimp fishery in the northwest Gulf of California. Note that although there is no overfishing condition (𝐻𝑅_MSY_ < 0.5), the ratio 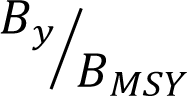 suggests signs of depletion but where the resource tends to recover.

## Discussion

Fishing is the only activity in the primary production sector where there are no inputs for production, and the exploitation of resources depends entirely on the natural biomass production capacity of populations. In a variable environment, this production capacity also varies, implying that management is supported by information on the available biomass. This is particularly relevant in annual species, such as the brown shrimp *P. californiensis*, where the success of the reproductive process is the only link between the population biomass in consecutive years; that is, the size of each annual cut will be a consequence of the recruitment caused by the reproductive stock of the previous year. Under these circumstances, it is very important, in management terms, to know both the available biomass and the key reference points that allow for dynamic sustainability of resource exploitation annually.

However, one of the major problems in many fisheries is the lack of data. In many fisheries, especially small-scale fisheries, there are no records of their operations; when information is available, it mainly corresponds to catch records or fishing effort records. However, for many resources, there is no consistent information on the structure of catches, and even less so in a broad temporal context that allows for detailed analysis of the dynamics of exploited populations. When time series information on catches and fishing effort records exists, dynamic biomass models are typically used [e.g., 35, 46, 47]. These models are based on the idea of partial survival, or remaining stock from one year to the next, which contributes to biomass production and the reproductive process. This does not occur in species with annual longevity, where the only link between successive years is the success of the reproductive process expressed as recruitment. In this context, in the present work, an analysis of the annual abundance decay of the brown shrimp population in the northwest Gulf of California (annual cohort) is developed based on monthly catch and fishing effort information.

In addition to the pattern of annual cohort abundance decay, the annual catchability coefficient was estimated, which, along with the catch per unit effort, allowed for monthly biomass estimates of the available population for all fishing seasons. These biomasses, related to the corresponding catches, allowed for an estimation of the harvest rate and, consequently, a first definition of the resource state of exploitation and the fishery’s evolution (Fig 6). Although this analysis suggests that the fishery activities are being carried out within the population’s surplus production limits, it does not provide information on management for future seasons; that is, it does not provide a prediction of available biomass to be taken as a criterion for decision-making.

In the developed analysis, two biological reference points were estimated: one objective, defined by the theoretical criterion of maximum sustainable yield, 𝐵𝑅𝑃_𝑡𝑔𝑡_, and a limit reference point, defined by the population replacement level 𝐵𝑅𝑃_𝑙𝑖𝑚_, i.e., the survival level of the adult population that, upon reproduction, gives rise to the recruitment level that exactly replaces the adult population. According to Fig 5, all fishing seasons were above the 𝐵𝑅𝑃_𝑙𝑖𝑚_, except for the years when the COVID-19 pandemic occurred, during which the fishery’s operation was entirely irregular, as shown by the fishing effort dynamics in Fig 1, particularly in the 𝑖𝑛𝑑𝑓 fleet, which accesses the adult phase of the population. In a precautionary context, if the confidence interval is considered when estimating 𝐵𝑅𝑃_𝑙𝑖𝑚_, the minimum risk level, for a survival rate 𝑠_𝑦_ = 0.13, corresponds to the upper limit of the recruitment rate given by the regression confidence interval, which corresponds to a recruitment rate 𝜌_𝑦_ = 0.38 and can be taken as the precautionary reference level for management.

Fig 8 shows the historical evolution of the harvest rates in contrast to 𝐵𝑅𝑃_𝑡𝑔𝑡_ and 𝐵𝑅𝑃_𝑙𝑖𝑚_. In all the cases, the harvest rates were below the different values of 𝐵𝑅𝑃𝑠. Fig 9 shows the equivalent conditions, but through the evolution of biomass changes. In the figure, the accumulated biomass at the end of the year, 𝐵_∞,𝑦_, the biomass corresponding to the maximum sustainable yield level, 𝐵_𝑀𝑆𝑌,𝑦_, the remaining biomass after exploitation, 𝐵_𝑟𝑒𝑚,𝑦_ = [𝐵_∞,𝑦_ ― (𝐻𝑅_𝑦_ ⋅ 𝐵_∞,𝑦_)], and the biomass corresponding to the 𝐵𝑅𝑃_𝑙𝑖𝑚_, showing the confidence interval values, are represented. As in the case of the harvest rates, the remaining population, which reflects the state of exploitation, was below the biomasses of 𝐵_∞,𝑦_ and 𝐵_𝑀𝑆𝑌,𝑦_ and above the biomasses corresponding to 𝐵𝑅𝑃_𝑙𝑖𝑚_. This suggests that this fishery operates at biologically sustainable production levels.

**Fig 8.**
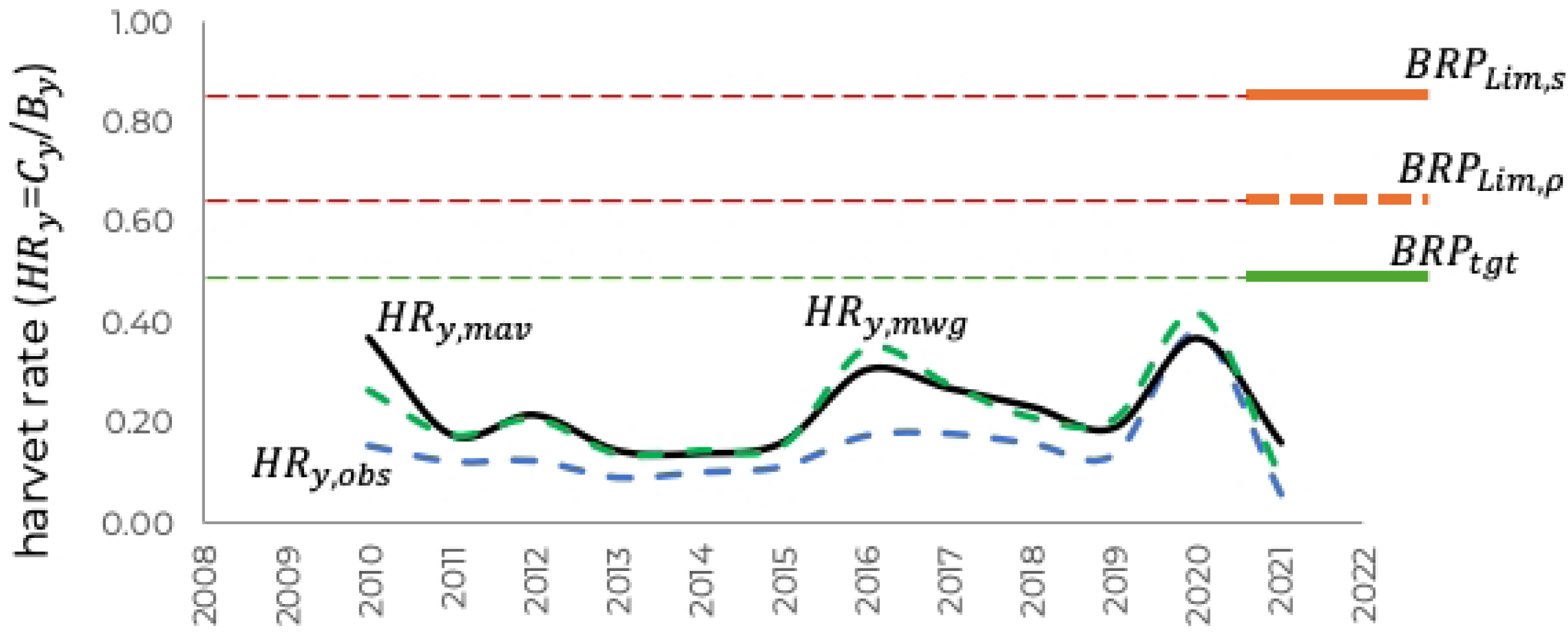
Annual trend of the estimated harvest rates for the target, 𝑩𝑹𝑷_𝒕𝒈𝒕_; and limit, 𝑩𝑹𝑷_𝒍𝒊𝒎,𝝆_(recruitment rate) and 𝑩𝑹𝑷_𝒍𝒊𝒎,𝒔_(survival rate), reference points; within a precautionary context.

**Fig 9.**
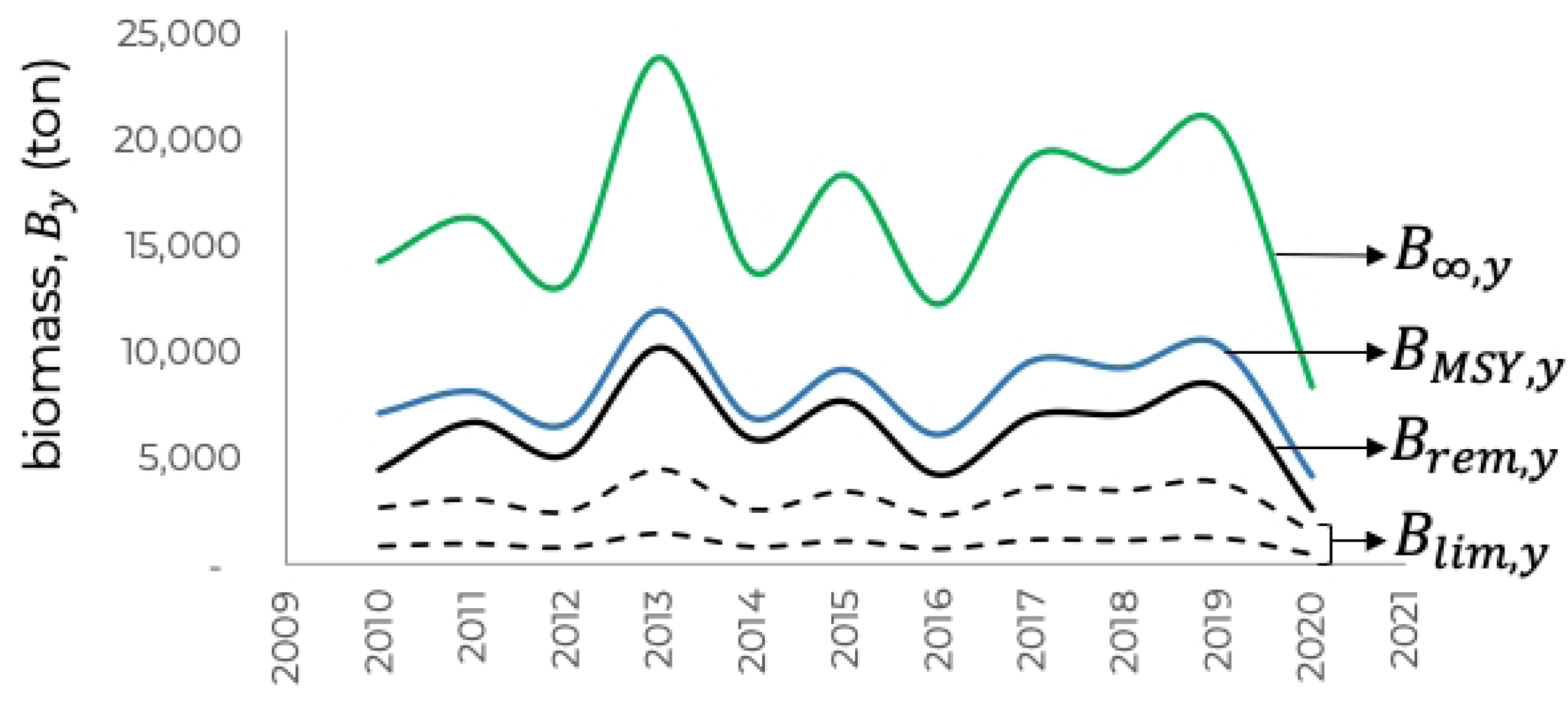
Annual trend of the biomass levels of the brown shrimp population associated with the. 𝑩𝑹𝑷𝒔: accumulated biomass in the year, 𝐵_∞,𝑦_; 𝐵_𝑀𝑆𝑌,𝑦_ as the biomass corresponding to the 𝐵𝑅𝑃_𝑡𝑔𝑡_; remaining biomass after fishing, 𝐵_𝑟𝑒𝑚_; and 𝐵_𝐿𝑖𝑚,𝑦_ as the biomass corresponding to the 𝐵𝑅𝑃_lim_ for each year, considering the limit recruitment and survival rates in a precautionary context. The remaining biomass was greater than the biomasses of the 𝐵𝑅𝑃_𝑙𝑖𝑚_ and less than the 𝐵𝑅𝑃_𝑡𝑔𝑡_.

Finally, the Kobe diagram represents the fishery trajectory, and the state of exploitation of the brown shrimp stock is shown in Fig 10, in which three reference levels are shown. The harvest rate 𝐻𝑅_𝑦_ = 0.60, which represents a level, albeit empirical, that is frequently assumed as a reference to the limit level in the absence of information about a fishery; this implies a population remnant of 40% after fishing. Other conditions of the fishery were also identified, the first referring to the limit reference point expressed by survival, 𝐵𝑅𝑃_𝑙𝑖𝑚,𝑠_, and the second corresponding to the limit reference point expressed by the minimum recruitment that guarantees the population’s replacement, 𝐵𝑅𝑃_𝑙𝑖𝑚,𝜌_.

**Fig 10.**
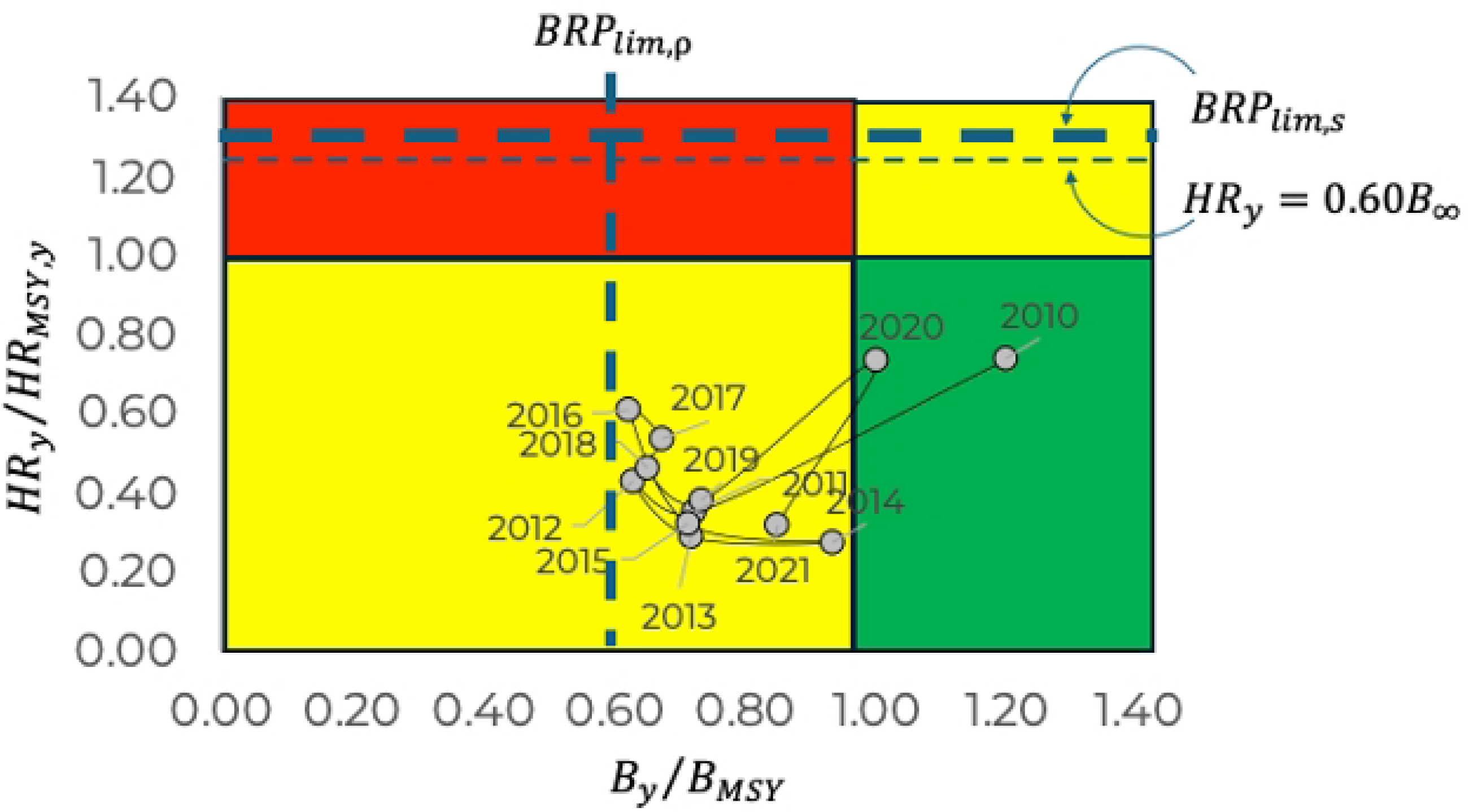
Kobe diagram for the *P. californiensis* brown shrimp fishery in the northwest Gulf of California, showing the state of the fishery in relation to different reference points. Note that the fishery operates at acceptable sustainability levels, indicated by the limit reference point that expresses the recruitment level required for population replacement under a precautionary criterion (minimum risk), and that, in terms of the fishing effect, it is far from critical levels.

The Kobe diagram in Fig 10, in contrast to that in Fig 7, illustrates two key aspects. First, over the past 20 years, the fishery has been operating under conditions that are acceptable for maintaining its biological sustainability. During the study period (2010 to 2021), the population level remained above the minimum recruitment level required for replacing the adult population 𝐵𝑅𝑃_𝑙𝑖𝑚,𝜌_. In contrast, the exploitation limits given by 𝐻𝑅_𝑦_ = 0.60 and the limit reference point represented by the survival, 𝐵𝑅𝑃_𝑙𝑖𝑚,𝑠_, are distant, in terms of the harvest rate, from the historical situation of the fishery.

The immediate conclusion is that the fishery is operating at acceptable sustainability levels, including interannual variability. However, it could be thought that it is operating very close to risk levels in relation to the recruitment level required for the replacement of the adult population. In this sense, it should be considered that this recruitment level corresponds to the minimum risk level, given the interannual variability of the study period, expressed by the upper level of the confidence interval in Fig 5. Importantly, the present analysis was based on catch and effort data, and the limit reference point shown in Fig 5 could be even lower if the analysis was based on the number of individuals; that is, considering that a unit of recruited biomass has a greater number of individuals than that same unit of biomass for adult organisms. This latter condition would probably result in a steeper slope of the relationship between survival rates and recruitment rates, leading to a lower biological limit reference point.

The current management strategy for this fishery, declared in the NFC (DOF 2018), is based on maintaining a minimum reproductive biomass at the end of the fishing season. To this end, surveys are conducted annually to evaluate the gonadal development status of the remaining population, defining the start of the closed season. This criterion fully corresponds to the 𝐵𝑅𝑃_𝑙𝑖𝑚,𝑠_(based on survival), so its definition can be useful for monitoring the fishery. Additionally, one of the management tactics mentioned in the NFC refers to the establishment of a spatiotemporal reproductive and growth ban. In the first case, it is the basis for maintaining reproductive biomass, as mentioned earlier; however, in the second case, although the document does not provide application details, the limit reference point linked to recruitment, 𝐵𝑅𝑃_𝑙𝑖𝑚,𝜌_, corresponds to the survival of recruits; therefore, in the sense of the growth ban, this reference point criterion could also be very useful.

Finally, the contrast of harvest rates against different reference criteria proves to be of great practical utility for monitoring the state of the fishery; for example, if a harvest rate is defined as a control criterion, being a function of the available biomass (𝐻𝑅 = 𝐶/𝐵), continuous review or change would not be necessary owing to environmental variations; that is, the feasible catch will always be a function of the available biomass regardless of biomass variability. The combination of the BRP limits based on the population replacement level, in combination with the definition and application of harvest rates, can be valuable elements for monitoring and controlling fishery sustainability.

## Acknowledgements

FAS and AHL thank the support of the EDI and COFAA programs of the Instituto Politécnico Nacional, as well as partial support through the SIP 20241264 and 20250571 projects. CIPQ thankful for the support of the Walton Family Foundation and International Community Foundation. Authors thank CONAPESCA for access to the data.

## Supporting information

**S 1. Parameters estimation for the relationships to compare fishing power between the SSF fleet and the IDF fleet.** The parameters were obtained using reduced major axis regression.

**S 2-A. Catch, fishing effort, catch per unit of effort, and available biomass for the 𝒔𝒔𝒇, per month and year for the brown shrimp, Penaeus californiensis, of the Gulf of California, Mexico.**

**S 2-B. Catch, fishing effort, catch per unit of effort, and available biomass for the 𝒊𝒏𝒅𝒇, per month and year for the brown shrimp**, *Penaeus californiensis*, **of the Gulf of California, Mexico.**

